# A microstructural analysis of water intake patterns associated with light phase food and water restriction in male and female rats

**DOI:** 10.64898/2026.11.03.764049

**Authors:** Emalie L. Mullane, Kaycee D. Moore, Andrea A. Edwards, Michael Tackenberg, Jessica Santollo

## Abstract

Water intake is influenced by both eating and circadian rhythms, and the available literature is mixed regarding which variable exerts greater control. Previous reports primarily focused on short term manipulations, however, one study using long term light phase food restriction found that water intake patterns primarily retained circadian characteristics in male rats. To further characterize this behavioral effect, water intake and lick microstructure was analyzed in male and female rats with food access restricted to the light phase. In addition, food and water intake and lick microstructure were analyzed when water access was restricted to the light phase (WRL). In support of the previous short term, but not long term, food manipulation studies, when food access was restricted to the light phase most water intake shifted to this time frame. This shift was medicated by increased lick burst number, suggesting altered satiety signaling underling this intake pattern. When water access was restricted to the light phase, food intake was equally distributed across the light and dark phases. There was no effect of sex on the patterns of intake during any test condition. These results further our understanding of the mechanisms underlying food and water-associated and circadian intake patterns by suggesting that altered satiety signaling underlies changes in water intake. Furthermore, these results show that the timing of food access has a greater effect on water intake patterns across the light-dark cycle than water access has on food intake patterns.

## Introduction

It is well known that most strains of laboratory rats are nocturnal and thus consume the majority of their food and water in the dark phase. This pattern of intake is observed as early as 21 days post-birth, once pups being feeding independent of the dam [1]. The percentage of dark phase intake does have a relatively large range with reported dark phase values between 60-95% [1–7], which may be related, in part, to strain differences [8, 9]. Regardless of this variability, it is well established that eating and drinking patterns are biological rhythms, controlled by circadian clocks. In support of this, rats housed under constant darkness continue to exhibit rhythms in their food and water intake patterns similar to those observed under light dark conditions [10, 11]. In addition, lesions to the suprachiasmatic nucleus, the central circadian clock, abolishes light-dark feeding and drinking patterns [12, 13].

In euhydrated rats, most water consumption is associated with food intake. Approximately 70% of water intake is consumed with food intake, primarily occurring within 15 minutes before or after a meal [6, 14]. Furthermore, when food intake is restricted or completely deprived, water intake decreases [15]. Water intake is, therefore, under the control of both food consumption and a circadian rhythm. Previous reports have attempted to disentangle these two variables to better understand the mechanisms underlying the rhythmic pattern of water intake. Short term (1 h) food access restricted to different times of the light phase increases light phase water intake, coinciding with the food access periods [1, 6, 16]. In addition, light phase palatable access has been reported to shift water intake into the light phase, even when standard chow was available in the dark phase [3]. A follow-up experiment, however, failed to replicate this effect [4]. In an attempt to reconcile these mixed findings, Spiteri [5] examined patterns of water intake across the light-dark cycle in male rats with food access restricted to the light phase for several weeks, to circumvent any interference that short term food presentation schedules may have on intake patterns. This report found that while some water intake shifted into the light phase, 76% of the daily water intake remained in the dark phase, leading to the conclusion that rats retain circadian characteristics in the temporal distribution of water intake independent of feeding.

Given the surprising retention of the circadian water intake pattern when food was restricted to the light phase, because rats are prandial drinkers, our primary goal was to further explore this behavioral pattern to better understand the circadian control of water intake. We, therefore, examined water intake in rats with food restricted to the light phase, but in addition to measuring total intake, a contact lickometer was used to allow for analysis of lick microstructure patterns. Lick microstructure analysis is a powerful tool because it allows consumption to be analyzed beyond total volume and provides insight into the physiological mechanisms underlying the ingestion [17–19]. In addition, because the previous studies were primarily conducted in males and there are well established sex differences in both ingestive behaviors and biological rhythms [20–27], we examined intake and probed for difference between the sexes as a secondary objective. Finally, to determine the effect light phase water access has on food intake patterns, we added an additional condition during which rats had water access restricted to the light phase with ab libitum food access throughout the day. The results from these studies will further our understanding of the mechanisms underlying food-associated and circadian related ingestive patterns in male and female rats.

## Methods

### Animals and Housing

Upon arrival into the facility, nine-and-a-half-week-old male and female Long Evans rats (n = 8/sex; Inotiv Laboratories, Indianapolis, IN) were singly housed in modified shoebox cages with *ad libitum* access to food (Teklad 2018; Harlan Laboratories) and tap water until otherwise noted. The temperature-and humidity-controlled colony rooms were maintained on a 12:12 h light-dark cycle (lights on at 0800 h). All experimental protocols were approved by the Animal Care and Use Committee at the University of Kentucky, and the handling and care of the animals was in accordance with the *National Institute of Health Guide for the Care and Use of Laboratory Animals*.

### Experimental Design

Data collection began after a one-week acclimation period. During the two-week baseline (BL) condition, rats had *ad libitum* access to food and water. The BL period was immediately followed by the food restricted to light phase (FRL) condition. During FRL, rats had food access restricted to the light phase (0800h – 2000h) with *ad libitum* access to water. After the FRL condition, rats had a one-week recovery period with *ad libitum* access to both food and water. The final condition of the experiment restricted water to the light phase (WRL). During WRL, rats had water access restricted to the light phase (0800h – 2000h) with *ad libitum* access to food. Daily body weights were recorded between 0700h – 0800h and food and water intakes were recorded as described below, but no intake measures were recorded during the one-week recovery period between the FRL and WRL conditions.

### Intake and Lick Microstructure Measures

Daily measures of water, food, and licks began at lights on (0800 h, ZT0) and concluded one hour prior to lights on (0700 h) to allow for 1 hour of daily animal maintenance (recording weights, refilling food/water, downloading lickometer data and restarting the computer). Food was weighed during the daily maintenance period (0700 h – 0800 h), 6 h into the light phase during weekdays (1400 h), and at dark onset (2000 h). Water bottles were weighed during the daily maintenance period and at dark onset. Daily licking behavior was measured using a contact lickometer (University of Kentucky, College of Arts and Sciences Electronics Shop) that recorded individual licks to allow for analysis of licking as a function of time and licking microstructure. The lickometer interfaced with a computer using an integrated USB digital I/O device (National Instruments, Austin, TX). Home cages were affixed with an electrically isolated metal plate with a 3.175-mm-wide opening, through which the rat needed to lick to reach the drinking spout, minimizing the possibility of non-tongue contact with the spout. A drinking (lick) burst was defined as at least 2 licks with an inter-lick-interval (ILI) of no more than 1 s and burst size was defined as the number of licks within a burst, as previously described [28–30].

### Data Analysis

All data are presented as means ± SEM throughout with significant p values determined at alpha < 0.05. Daily intakes were averaged during the BL condition for each animal. Five of the baseline days were excluded from analysis (resulting in 9 baseline days used in the analysis) due to high humidity in the colony room (> 2 standard deviations above the mean) during the first 3 days of the BL period. In addition, on BL Day 9 there was a power outage in the animal facility which extended the dark phase for an addition 3 hours. We, therefore, excluded all data from this day and the following day (BL Day 10). For the FRL and WRL measures, the daily percentage of light phase consumption was analyzed and compared to the average BL value to determine when intake became stable. Then, the daily intakes were averaged across the stable portion of the FRL and WRL periods and used for the subsequent analyses. Technical issues with the lickometer recordings resulted in the loss of 1-3 daily recordings in 6 rats during the FRL condition and 2-3 daily recordings in 5 rats during the WRL condition. T-tests were used to determine the effect of sex on the percent light phase consumption during baseline. All additional analysis used a mixed design ANOVA with sex as the between-subjects factor and light phase, testing condition (BL, FRL and/or WRL), and/or time (hourly bin) as the within-subjects factors. To estimate the phase of the licking behavior, mean licks per hour across BL and FRL were fit using a fixed-frequency linear model *E*(*t*) = *Β*_0_ + *β*_1_*xs* + *β*_2_*xc*, where 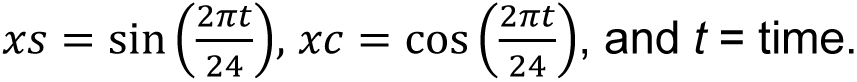 The phase of the peak of the fitted wave was calculated as 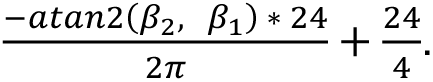 Circular mean, the mean resultant vector length (rho), and circular ANOVA were calculated using the circular R package [31]. Newman-Keuls pairwise comparisons were used to follow up main or interactive effects.

## Results

### Intake patterns during BL and stabilization of intake and body weight during FRL and WRL

As expected, during the BL period all intake measures were greater in the dark phase compared to the light phase. Statistical analyses of these patterns are incorporated into the analysis of intake during the FRL and WRL conditions below. Rats consumed 35.53% of their food during the light phase and this did not differ by sex, t(14) = 0.25, p = n.s. Similarly, rats consumed 35.00% of their water during the light phase and this did not differ by sex, t(14) = 0.51, p = n.s.

During FRL, the percent of water consumed during the light phase increased compared to the BL period. Analysis of the daily percent of light phase water consumption revealed a main effect of day, F(14,196) = 33.60, p < 0.001, but not sex, F(1,14) = 1.55, p = n.s., nor an interaction between sex and day, F(14,196) = 0.86, p = n.s. (Figure 1). Post hoc analysis revealed that the percent of water consumed during the light phase from FRL Day 5-14 was stable [69.3-76.7%], with no statistical difference between these days. Percent light phase intake on FRL Days 5-14 was greater than during BL and FRL Days 1 and 2. During WRL, the percent of food consumed during the light phase increased compared to the BL period. Analysis of the daily percent of light phase food consumption revealed a main effect of day, F(14,196) = 6.27, p < 0.001, but not sex, F(1,12) = 0.15, p = n.s., nor an interaction between sex and day, F(14,196) = 0.86, p = n.s. (Figure 1). Post hoc analysis revealed that the percent of food consumed during the light phase from WRL Day 4-14 was stable [41.4-49.8%], with no statistical difference between any of these days (except for WRL Day 9 > WRL Day 6). Percent light phase intake on WRL Days 4-14 was greater than during BL and WRL Day 1. Subsequent analysis of intake during the testing conditions were restricted to the average values from FRL Days 5-14 and WRL Days 4-14.

**Figure 1.**
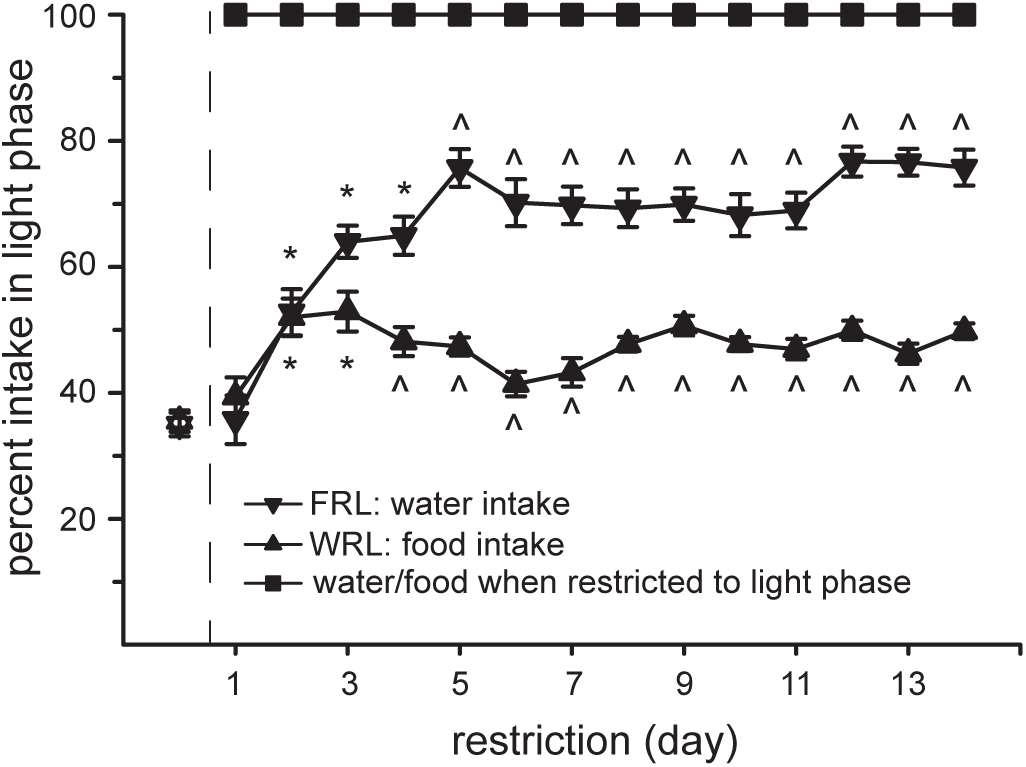
Food or water restriction to the light phase increased light phase water or food intake, respectively. During ab libitum food and water access, about 35% of intake occurred in the light phase (open symbols). When food was restricted to the light phase, water intake in the light phase increased to 72.1%, which stabilized on Day 5. When water was restricted to the light phase, food intake in the light phase increased to 47.2%, which stabilized on Day 4. Vertical dashed line indicates the start of the restriction periods. *Greater than baseline, p < 0.05. ^Greater than baseline and not different from Day 14.

Weekly body weights were influenced by a main effect of sex, F(1,14) = 163.17, p < 0.001, week, F(7,98) = 208.55, p < 0.001, and an interaction between sex and week, F(7,98) = 57.76, p < 0.001 (Table 1). As expected, males weighed more than aged-matched females, p < 0.001. In males, body weight generally increased week to week, however, one week after the start of FRL (Week 4), body weights were significantly less than the previous week. At the end of the FRL condition (Week 5), male body weights returned to where they were at on the start of Week 3. The WRL condition had no effect on weekly weight gain in males. In females, body weight generally increased week to week, similar to the males. Again, there was a decrease in body weight after one week of FRL. At the end of the FRL condition, body weight increased and was not significantly different from weights at either the start of Week 3 or Week 4. Unlike the males, after one week of WRL (Week 7) there was no significant increase in body weight in females, as weights remained statistically similar to those at the start of the WRL condition (Week 6). By the conclusion of the WRL condition, body weights in the females were greater than at the start of the WRL phase (Week 6).

**Table 1.** Body Weights at the start of each week.

|  | Baseline |  | FRL |  | Off Week | WRL |  |  |
|---|---|---|---|---|---|---|---|---|
|  | Week 1 | Week 2 | Week 3 | Week 4 | Week 5 | Week 6 | Week 7 | End |
| Males | 288.05 ± 6.33<sup>a</sup> | 334.38 ± 8.27<sup>b</sup> | 369.85 ± 10.51<sup>c</sup> | 355.30 ± 10.59<sup>d</sup> | 368.88 ± 10.69<sup>c</sup> | 402.04 ± 12.77<sup>e</sup> | 421.13 ± 14.27<sup>f</sup> | 446.25 ± 15.68<sup>g</sup> |
| Females | 200.99 ± 2.85<sup>h</sup> | 217.88 ± 3.36<sup>i</sup> | 229.99 ± 2.94<sup>j</sup> | 218.33 ± 3.26<sup>i</sup> | 221.61 ± 3.22<sup>i,j</sup> | 234.96 ± 3.85<sup>k</sup> | 242.71 ± 5.24<sup>k,l</sup> | 250.99 ± 6.13<sup>l</sup> |

Different letters denote differences between different conditions.

### Total intake and lick microstructure across testing conditions

Total food intake was influenced by a main effect of sex, F(1,14) = 133.79, p < 0.001, and testing condition, F(2,28) = 40.34, p < 0.001, but not an interaction of sex and testing condition, F(2,28) = 1.25, p = n.s. (Figure 2A). Males consumed more food than females, p < 0.001. Food intake was greatest during BL, lowest during FRL, with an intermediate level of food intake during WRL, p < 0.01. Total water intake was influenced by a main effect of sex, F(1,14) = 52.73, p < 0.001, and testing condition, F(2,28) = 74.08, p < 0.001, but no interaction of sex and testing condition, F(2,28) = 1.29, p = n.s. (Figure 2B). Males consumed more water than females, p < 0.001. Total water intake was reduced during WRL compared to BL and FRL, p < 0.001. Analysis of total licks revealed the same result, a main effect of sex, F(1,14) = 5.84, p < 0.05, and testing condition, F(2,28) = 103.51, p < 0.001, but no interaction between sex and testing condition, F(2,28) = 0.78, p = n.s. (Figure 2C). Males had more licks for water than females, p < 0.001. Total licks were reduced during WRL compared to BL and FRL, p < 0.001. Total burst number was influenced by a main effect of testing condition, F(2,28) = 42.75, p < 0.001, but not sex, F(1,14) = 2.63, p = n.s., nor an interaction between sex and testing condition, F(2,28) = 1.28, p = n.s. (Figure 2D). While burst number increased during FRL compared to BL, it decreased during WRL compared to BL, p< 0.001. Finally, burst size was influenced by a main effect of sex, F(1,14) = 11.13, p < 0.005, and testing condition, F(2,28) = 11.74, p < 0.001, but no interaction of sex and testing condition, F(2,28) = 2.15, p = n.s. (Figure 2E). Males had larger burst sizes compared to females, p < 0.005. Burst sizes during FRL and WRL were smaller than BL, p < 0.001.

**Figure 2.**
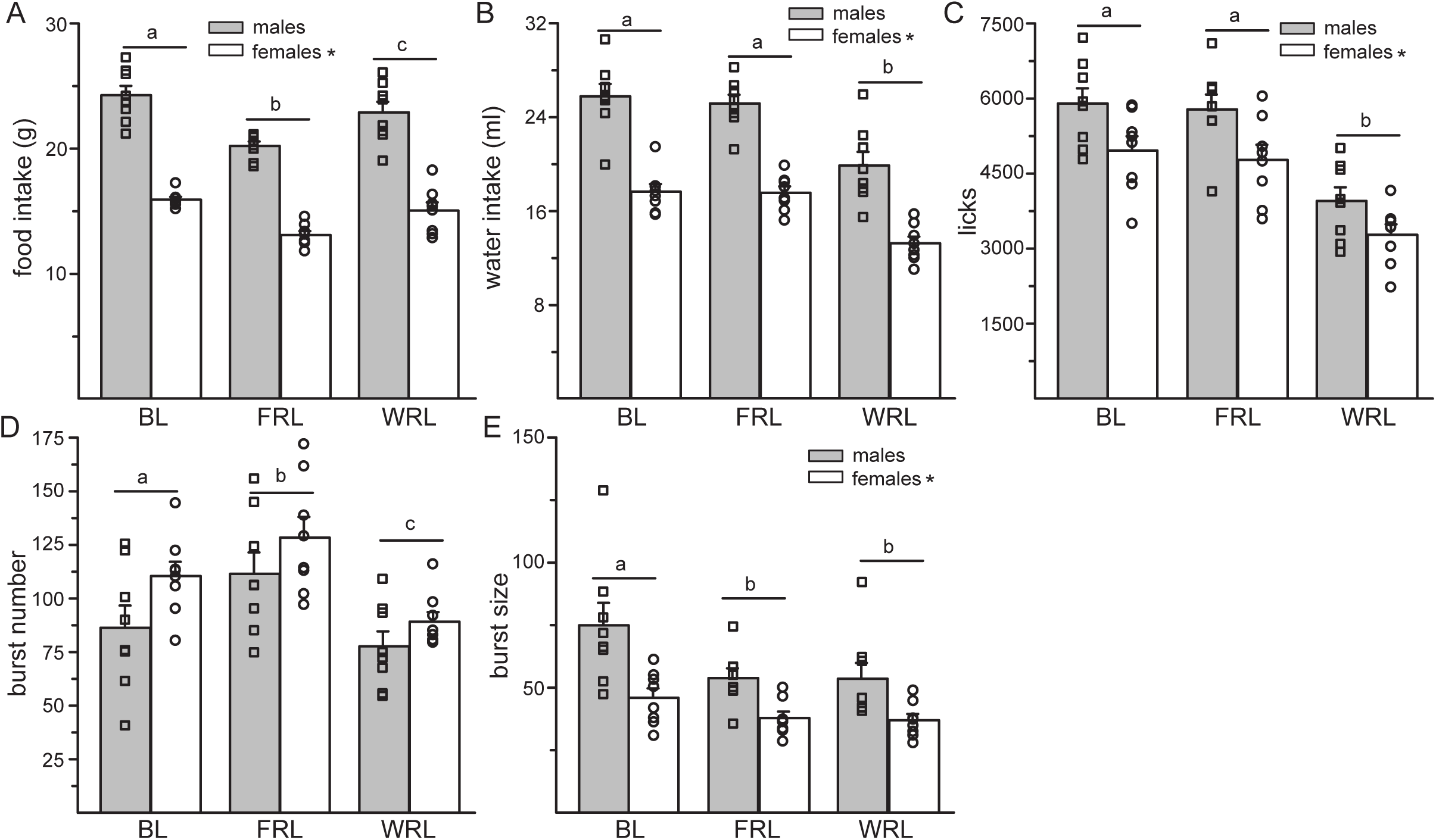
Limiting food or water access to the light phase influenced total intake. Males consumed more, took more licks, and had larger burst sizes than females. There was no effect of sex on total burst number (A-E). (A) During both FRL and WRL, total food intake was reduced compared to the BL period, with total food intake during the FRL being less than both BL and WRL. (B/C) Total water and licks were reduced, compared to BL, during the WRL. There was no change in total water or licks during FRL. (D) Compared to BL, FRL increased total burst number while WRL decreased total burst number. (E) Both FRL and WRL reduced total burst size, compared to BL. *Less than males, p < 0.05. Different letters denote differences between different experimental conditions.

### Intake and microstructure patterns during FRL

Water intake during FRL was influenced by sex, F(1,14) = 71.66, p < 0.001, and an interaction between testing condition and light phase, F(1,14) = 296.11, p < 0.001, but no other main or interactive effects, F(1,14) = 0.001-3.33, p = n.s. (Figure 3A). Males consumed more water than females, p < 0.001. During BL more water was consumed in the dark, compared to the light phase. During FRL, however, more water was consumed during the light phase than the dark phase. Furthermore, while BL dark phase and FRL light phase water intakes were similar, intake during the FRL dark phase was less than all other conditions, p < 0.05.

**Figure 3.**
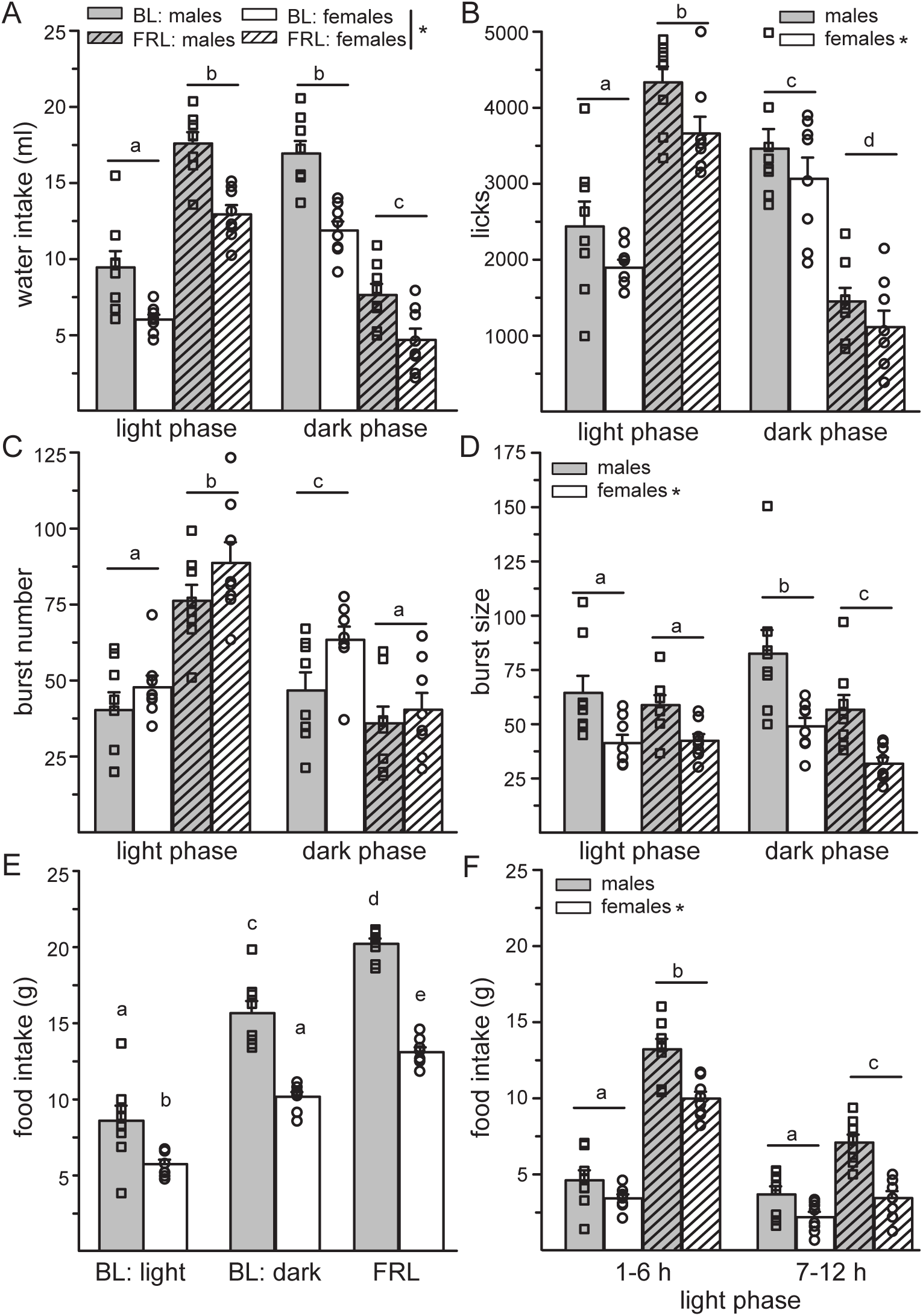
Restricting food to the light phase changed water intake patterns. Males consumed more, took more licks, and had larger burst sizes than females. There was no effect of sex on burst number (A-F). (A) During BL, water intake was greater in the dark phase, compared to the light phase. During FRL, water intake was greater in the light phase, compared to the dark phase. Dark phase intake during FRL was less than all other times. (B) A similar finding occurred for licks, however, FRL light phase licks were even greater than BL dark phase licks. (C) During BL, burst number was greater in the dark phase, compared to the light phase. During FRL, burst number was greater in the light phase, compared to the dark phase, and greater than during the BL dark phase. (D) During BL, burst size was increased in the dark phase, compared to the light phase. During FRL, burst size was decreased in the dark phase, compared to the light phase. During the light phase, burst size was similar between BL and FRL. (E) Twelve-hour food intake was greatest during FRL, with BL dark phase intake greater than BL light phase intake. (F) During FRL, more food was consumed in the first half of the light phase, compared to the second half. During BL, there was no difference in food intake between the two halves of the light phase. *Less than males, p < 0.05. Different letters denote differences between different experimental conditions.

Licks for water during FRL was influenced by sex, F(1,14) = 5.90, p < 0.05, light phase, F(1,14) = 18.18, p < 0.001, and an interaction between testing condition and light phase, F(1,14) = 131.90, p < 0.001, but no other main or interactive effects, F(1,14) = 0.07-1.41, p = n.s. (Figure 3B). Males had more licks for water than females, p < 0.05. Unlike during BL when more licks occurred in the dark phase, during FRL more licks for water were taken during the light phase. Furthermore, licks during the FRL light phase were greater than licks during the BL dark phase. Licks during the FRL dark phase were less than all other conditions, p < 0.005. This shift in water intake was driven by changes in burst number. Burst number was influenced by a main effect of testing condition, F(1,14) = 32.50, p < 0.001, light phase, F(1,14) = 34.50, p < 0.001, and an interaction between testing condition and light phase, F(1,14) = 90.59, p < 0.001, but no other main or interactive effects, F(1,14) = 0.01-2.67, p = n.s. (Figure 3C). While the number of bursts was greater in the dark phase during BL, during FRL burst number was greater in the light phase, p < 0.001. Furthermore, burst number during the FRL light phase was greater than BL dark phase burst number, p < 0.001. Burst size was influenced by a main effect of sex, F(1,14) = 10.85, p < 0.01, testing condition, F(1,14) = 25.64, p < 0.001, and an interaction between testing condition and light phase, F(1,14) = 63.39, p < 0.001, but no other main or interactive effects, F(1,14) = 0.07-3.34, p = n.s. (Figure 3D). Males had larger burst sizes than females, p < 0.01. During BL, burst size was greater during the dark phase, compared to the light phase. During FRL, burst size was greater during the light phase, compared to the dark phase. FRL light phase burst size, however, was not different from BL light phase burst size and was smaller than BL dark phase burst size, demonstrating that there was no change in burst size that mimicked the increased water intake during this time, p < 0.001.

Analysis of food intake in 12 h bins (BL light phase, BL dark phase, FRL) revealed a main effect of sex, F(1,14) = 208.29, p < 0.001, a main effect of 12 h bin, F(2,28) = 111.95, p < 0.001, and an interaction between sex and 12 h bin, F(2,28) = 5.67, p < 0.01 (Figure 3E). In both males and females, food intake was greatest during the FRL, and food intake was greater in the BL dark phase compared to the BL light phase, p < 0.001. Males consumed more food than females, p < 0.001, with male intake during the BL light phase being comparable to female intake during the BL dark phase. Finally, analysis of food intake in 6 h bins revealed that the majority of intake during FRL occurred early in the light phase. Food intake was influenced by a main effect of sex, F(1,14) = 51.34, p < 0.001, testing condition, F(1,14) = 455.91, p < 0.001, time, F(1,14) = 72.19, p < 0.001, and an interaction between testing condition and time, F(1,14) = 44.52, p < 0.001 (Figure 3F). Again, males consumed more food than females and more food was consumed during the light phase under the FRL condition, p < 0.001. During BL, there was no difference in food intake between the first and second half of the light phase. During the FRL, however, more food was consumed during the first half of the light phase, compared to the second half. Intake in both halves of the light phase was greater during the FRL condition compared to BL, p < 0.001.

Analysis of hourly licks during BL revealed a main effect of sex, F(1,14) = 5.21, p < 0.05 and a main effect of time, F(22,308) = 15.70, p < 0.001 but no interaction between sex and time, F(22,308) = 0.98, p = n.s. (Figure 4A). Males had more licks for water than females, p < 0.05. Licks for water were greatest during hours 1 and 12 (light phase) and 23 (dark phase). Licks during hours 14-22 (dark phase) were greater than licks during hours 2-11 (light phase). Waveform fits of mean licks per hour during BL featured a peak at 21.02 h for males and 19.27 h for females, F(1, 14) = 3.628, p = n.s. (Figure 4B). Analysis of hourly licks during FRL revealed a main effect of sex, F(1,14) = 5.64, p < 0.05, a main effect of time, F(22,308) = 139.52, p < 0.001, and an interaction between sex and time, F(22,308) = 2.49, p < 0.05 (Figure 4C). Males had more licks for water than females, p < 0.05. Licks for water were greatest during hour 1 (light phase), licks during hour 2 (light phase) were greater than during hours 3-23 and licks during hour 6 (light phase) were greater than during hours 3-5 and 7-23. Similar intake patterns were observed between males and females except that the increased intake during hour 2, compared to hour 6, was driven by the females, p < 0.05. Waveform fits of mean licks per hour during FRL featured a peak at 2.24 h for males and 1.83 h for females, F(1, 14) = 0.828, p = n.s. (Figure 4D).

**Figure 4.**
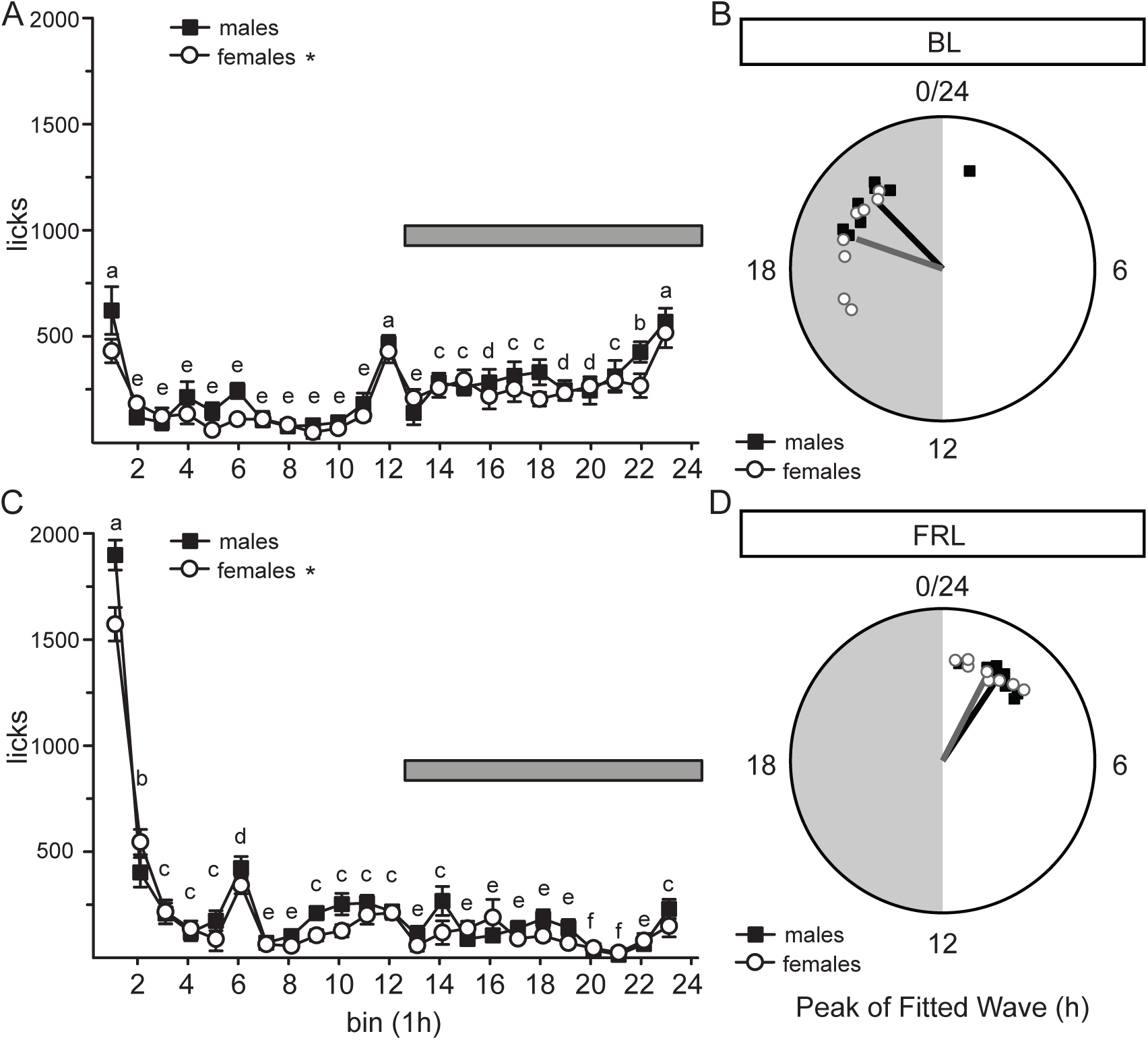
Licks for water across time. (A) Licks for water in 1-hour bins during BL. Regardless of hour, males had more licks for water than females. Licks for water were greatest during hours 1, 11, and 23. Licks for water during hours 13-22 (dark phase) were greater than licks for water during hours 2-10 (light phase). (B). Peak phase of a fitted waveform model to the mean hourly lick values during BL for males (closed squares) and females (open circles). The angle of the radial lines represent the circular mean of the groups, and the length of the line represents the mean resultant vector length (rho) (C) Licks for water in 1-hour bins during FRL. Regardless of hour, males had more licks for water than females. Licks for water were greater during hour 1, licks for water during hour 2 were greater than hours 3-23, and licks for water during hour 6 were greater than hours 3-5 and 7-23. (D) Peak phase of a fitted waveform model to the mean hourly lick values during FRL for males (closed squares) and females (open circles). The angle of the radial lines represent the circular mean of the groups, and the length of the line represents the mean resultant vector length (rho). *Less than males, p < 0.05. Different letters denote differences between different hours. The dark phase is denoted with a horizontal bar (A, C) or grey shading (B, D).

**Figure 5.**
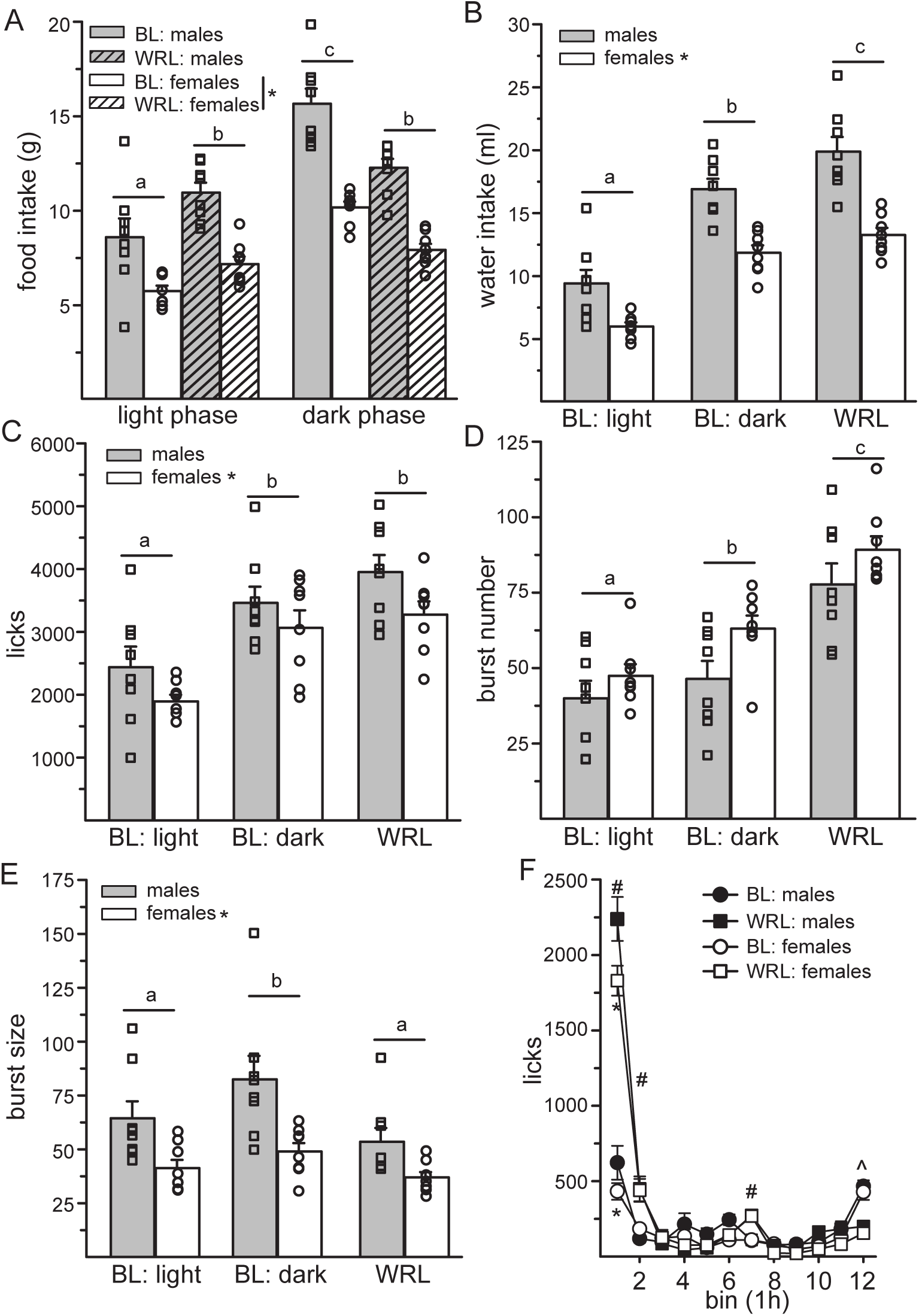
Restricting water to the light phase changed food intake patterns and lick microstructure. Males consumed more, took more licks, and had larger burst sizes than females. There was no effect of sex on burst number (A-F). (A) During BL, food intake was greater in the dark phase, compared to the light phase. During WRL, food intake was similar between the light and dark phase and, compared to BL, it was in between intake levels in the light and dark phases. (B) Twelve-hour water intake was greatest during WRL, with BL dark phase intake greater than BL light phase intake. (C) Twelve-hour licks during WRL were greater than BL light phase, but not different from BL dark phase. (D) Twelve-hour burst number was greatest during WRL, with BL dark phase burst number greater than BL light phase burst number. (E) Twelve-hour burst size during WRL was similar to BL light phase which were both smaller than BL dark phase burst size. (F) Light phase licks were greater during WRL than BL. During the first hour of intake, males had more licks than females. During hours 1, 2, and 7, more licks were taken during WRL than BL. During hour 12, more licks were taken during BL than WRL. *Less than males, p < 0.05. #Greater than BL, p < 0.05. ^Greater than WRL, p < 0.05. Different letters denote differences between different experimental conditions.

### Intake and microstructure patterns during WRL

Food intake during WRL was influenced by sex, F(1,14) = 82.79, p < 0.001, testing condition, F(1,14) = 11.23, p < 0.005, light phase, F(1,14) = 47.56, p < 0.001 and an interaction between testing condition and light phase, F(1,14) = 32.63, p < 0.001, but no other main or interactive effects, F(1,14) = 0.14-2.65, p = n.s. (Figure 4A). Males consumed more food than females, p < 0.001. Unlike during BL when more food was consumed during the dark phase, during FRL rats consumed similar amounts of food between the light and dark phases, an amount of which was greater than BL light phase but less than BL dark phase, p < 0.001. Analysis of FRL light phase food intake in 6 h bins found a main effect of sex, F(1,14) = 36.43, p < 0.001 and bin, F(1,14) = 107.88, p < 0.001, but no interaction between sex and bin, F(1,14) = 0.22, p = n.s. More food was consumed in the first half of the light phase (6.12 ± 0.36), compared to the second half (3.11 ± 0.30; data not shown).

Water intake during WRL was influenced by sex, F(1,14) = 39.70, p < 0.001 and 12 h bin, F(2,28) = 84.21, p < 0.001, but no interaction between sex and 12 h bin, F(2,28) = 2.52, p = n.s. (Figure 4B). Water intake was greater in males, compared to females, p < 0.001. Water intake was greatest during WRL, with water intake during the BL dark phase greater than during the BL light phase, p < 0.001. Licks for water during WRL was influenced by sex, F(1,14) = 5.05, p < 0.05 and 12 h bin, F(2,28) = 22.04, p < 0.001, but no interaction between sex and 12 h bin, F(2,28) = 0.19, p = n.s. (Figure 4C). Licks for water were greater in males, compared to females, p < 0.05. Licks during the BL light phase were less than during the BL dark phase and the WRL, p < 0.001. Again, the increased water intake during WRL was mediated by a change in burst number. Burst number was influenced by a main effect of 12 h bin, F(2,28) = 82.21, p < 0.001 but not sex, F(1,14) = 3.26, p = n.s. nor an interaction between sex and 12 h bin, F(2,28) = 1.04, p = n.s. Burst number was greatest during WRL, with burst number during the BL dark phase greater than during the BL light phase, p < 0.005. Burst size was influenced by a main effect of sex, F(1,14) = 9.63, p < 0.01 and 12 h bin, F(2,28) = 12.19, p < 0.001 but no interaction between sex and 12 h bin, F(2,28) = 2.08, p = n.s. (Figure 4E). Males had larger burst sizes than females, p < 0.01. Burst size during the BL dark phase was larger than burst size during both the BL light phase and WRL, p < 0.005.

Hourly licks during the light phase were influenced by a main effect of testing condition, F(1,14) = 122.55, p < 0.001, time, F(11,154) = 253.52, p < 0.001, an interaction between sex and time, F(11,154) = 4.10, p < 0.001, and an interaction between testing condition and time, F(11,154) = 103.33, p < 0.001, with no other main or interactive effects, F(1/11,14/154) = 0.26-3.70, p = n.s. (Figure 4F). Males had more licks for water compared to females during the first hour of the light phase. Licks for water during hours 1, 2, and 7 were greater during WRL, compared to BL. Licks for water during hour 12 were greater during BL, compared to WRL, p < 0.001.

### Comparison of the change in light phase consumption between testing conditions

During FRL, significantly more water intake was consumed in the light phase, compared to the dark phase (Figure 3A). Durling WRL, food intake was roughly equivalent between the light and dark phases (Figure 4A). To statistically demonstrate that light phase food restriction had a greater effect on water intake than light phase water restriction had on food intake, the percent change from BL in light phase consumption between water intake during FRL and food intake during WRL was analyzed. There was a main effect of condition, F(1,14) = 133.79, p < 0.001 but no effect of sex, F(1,14) = 0.01, p = n.s. nor interaction between condition and sex, F(1,14) = 4.57, p = n.s. (Figure 1). Restricting food to the light phase had a greater effect on light phase water intake than restricting water to the light phase had on food intake during the light phase, p < 0.001.

## Discussion

The goal of these experiments was to characterize the lick microstructure patterns, and probe for sex differences, that occur when either food or water access is restricted to the light phase. The results demonstrate that when food access was restricted to the light phase, the majority of water intake shifted to this time, resulting in a 37-percentage point increase in the percentage of light phase water consumption (35% to 72%). This shift in water intake was the result of an increase in burst number during the light phase with no change in burst size during this time. When water access was restricted to the light phase, food intake shifted into the light phase, however, at a reduced magnitude in comparison to the shift in water intake. Food intake was roughly equally distributed between the light and dark phases, resulting in a 12-percentage point increase in the percentage of light phase food consumption (35% to 47%). Limiting water access to the light phase, again, was driven by an increase in burst number, with no change in burst size. These findings are in agreement with four previous reports demonstrating changes in water intake patterns due to alterations in food access [1, 3, 6, 16], but in disagreement with two others which reported a maintenance of the circadian pattern of water intake [4, 5]. These results also show no effect of sex on the changes in intake patterns under these manipulations, an important variable to understand given sex differences in ingestive behaviors and biological rhythms [20–27]. Together these results suggest that the timing of feeding can disrupt the circadian pattern of water intake and, while water intake also influences food intake patterns, it appears to be less pronounced than the effect eating has on drinking patterns.

During the two-week baseline period, as expected, all measures of ingestive behaviors (food and water intake, licks, burst number and burst size) were greater during the dark phase than the light phase. Sixty-five percent of both food and water intake occurred in the dark phase, which falls within previously reported ranges of dark phase consumption (60-90%), albeit at the lower end of said range. Reduced dark phase intake could have been influenced by our intake measurements. Because the experiment’s manipulation was to restrict food or water access to the light phase, dark phase consumption was limited to 11 hours to allow for 1 h of daily animal maintenance. Furthermore, food was manually measured 6 h into the light phase and food and water were manually measured at the end of the light phase which likely aroused the animals. Together these factors likely biased some intake, or intake percentage, towards the light phase. In support of this theory, Siegel [2] measured daily food and water intake in 6 h intervals and observed dark phase intake (70%) on the lower end of the range reported in the literature. Hourly water intake during the BL period was greatest during the first and last h of the light phase and the last hour of dark phase access. The increased water intake at the end of the dark phase corroborates previous reports in rats and mice [5, 32, 33] which is hypothesized to protect against dehydration during sleep [32]. The high levels of water intake during the first hour of the light phase has previously been reported in Long-Evans rats [8], but could also be the result of a shift in the timing of this daily anticipatory water intake [32] since rats lost access to the water bottles during the last h of the dark phase (for daily maintenance). There was no effect of sex on light phase patterns of water or food intake or hourly licks, although males did have greater food and water intake, more licks for water and larger burst sizes. We recently demonstrated a sex difference in burst size, but not burst number, during dehydration induced water intake [30]. The findings here, therefore, extend this previous finding to non-emergency, or endogenous [17], water intake, together revealing that under multiple conditions of water intake burst size is greater in males than females.

A primary goal of this series of experiments was to further characterize water intake patterns via examining the lick microstructure when food access was restricted to the light phase in male and female rats. In contrast to the report by Spiteri [5], the present results show that the pattern of water intake and licks reversed, with most water intake occurring during the light phase, in particular the first hour of the light phase, when rats had access to food. Of note, the licking behavior does not appear to undergo a complete inversion during FRL, as waveform fits of mean hourly lick data in FRL (Figure 4D) show a roughly 6-hour delay in peak time relative to BL (Figure 4B), rather than a 12-hour change. Peak times in BL occurred primarily in the second half of the dark phase and peak times in FRL occurred in the first half of the light phase, though the fits during FRL were likely influenced by the high lick values recorded during the first hour (Figure 1C). The Spiteri study also reported a decrease in total daily water intake, compared to the baseline period, however total daily water intake and licks during the FRL were not different than during BL in the present study. Lick microstructure analysis revealed that the shift in water intake in the light phase was driven by an increase in burst number suggesting altered satiety signals associated with water intake under this condition [18, 19, 34]. There was no change in light phase burst size between the BL and FRL conditions, although dark phase burst size decreased in the absence of food. Although our food intake measurements were less refined than Spiteri, the results are similar to those previously reported. When restricted to the light phase, total food intake was decreased, compared to BL, and most of the food intake occurred in the first half of this time period. Spiteri [5] reported that 50% of the food intake during FRL was in the first h of the light phase and while our food measurement were only in 6 h bins, the majority of the licks for water were taken during the first h of the light phase, suggesting that food intake was likely concentrated around this time. Finally, the changes in water and food intake patterns did not differ between males and females, suggesting no sex difference in these ingestive behavior patterns.

Our results do not support the findings of Spiteri [5], which inspired this series of experiments, but they are in agreement with other studies which observed changes in water intake patterns in response to short-term light phase food restriction [1, 3, 6, 16]. Furthermore, despite the maintenance of the circadian characteristics in the temporal distribution of water intake in the Spiteri and Greenwood reports [4, 5], the distribution of light to dark phase water intake dampened in these studies with some water intake (9-19 percentage points) shifting into the light phase. It is difficult to reconcile the discrepancies between these studies without further research. Spiteri’s study was designed to circumvent any interference presented by the short food presentation schedules and hence monitored intake patterns across four weeks of light phase food restriction. While our study limited this period to only two weeks, we found stable intake patterns by Day 5 and Spiteri reported the maintenance of dark phase water intake across all 4 weeks. The percentage of dark phase consumption during the BL periods were different between the two studies, with 95% dark phase intake in the Spiteri study vs 65% in the current study. This could contribute to the different outcomes, as the threshold to flip the majority water intake into the light phase was lower for our study. However, the magnitude of the change in intake was still greater in our study (37 percentage point change in light phase water intake) compared to Spiteri (19 percentage point change), therefore we do not believe the different baseline values are the cause of the discrepancy. A limitation may be that the baseline dark phase intakes are at both extremes (65% and 95%) of the reported ranges (60-95%) in our study and that of Spiteri, respectively [1–7]. Baseline dark phase water intake in the short access studies ranged from 65% [3] to 87% [35], and again these reports found that water intake shifted to concentrate around the light phase food access. We, therefore, do not believe the discrepancy in these results is due to the baseline light to dark phase distributions. Furthermore, because we detected no sex difference in intake patterns in the present study, it is unlikely that the sex of the animals contributed to these discrepancies. Strain differences could be a source of discrepancy, as differences in circadian feeding and drinking patterns between strains exist [8, 9], though differences in food associated water intake patterns have also been reported within a single strain [5, 6]. The source of the discrepancies between these studies remains unclear and may be some combination of small differences in strain, testing timelines, and baseline light to dark phase intake distributions.

An additional goal of this series of experiments was to determine how restricting water intake to the light phase influences food intake patterns while probing for sex differences. Total food intake decreased, compared to BL, although the magnitude of this reduction was less than when food access was restricted to the light phase. Food intake became roughly equally distributed across the light-dark phases with 47% of intake occurring in the light phase, with about twice as much food eaten in the first half of the light phase compared to the second half. Total water intake and licks decreased compared to BL when access was restricted to the light phase. Most of the water intake occurred during the first 2 h of light phase water access, with a small but significant increase in licks in the middle of the light phase. Only during the last h of the light phase was water intake significantly reduced in the WRL condition, compared to BL. While total burst number and burst size decreased, mimicking the reduction in overall water intake, light phase burst number during WRL increased compared to burst number in both the BL light and dark phases. During WRL, burst size did not increase and remained similar in size to bursts taken during the BL light phase. Again, the changes in water and food intake patterns did not differ between males and females, further supporting the null effect of sex on water and food intake patterns.

Lick microstructure analysis allows for water intake to be analyzed beyond overall volume and provides insight into the hedonic and satiating properties underling the consumption. Water, and other fluid, intake patterns across a test session occur in bursts, often defined as at least two licks with an interlick interval of less than 1000 ms and common lick microstructure analyses quantify the number of bursts and the average size of the licking bursts across a test session. Research by Davis and Smith demonstrated that the burst number associated with increasing concentrations of sucrose follows an inverted U-shaped curve and that sham feeding paradigms result in increased burst number [36–38]. Furthermore, increases in water intake in water-deprived rats under sham-drinking conditions are primarily driven by changes in burst number [34]. Together, these studies led to the conclusion that changes in burst number represent changes in satiety feedback signals. Changes in burst size, on the other hand, follows the pattern of intake when the concentration of sucrose is manipulated or adulterated with quinine [37–40]. In addition, recent studies demonstrate that increased intake of cold water, which is preferred over tepid water, by water-deprived rats, is driven by changes in burst size [30]. Together, these findings led to the conclusion that burst size represents changes in the hedonic value of the fluid. When food was restricted to the light phase, the majority of water intake shifted to this time. This shift was driven by an increase in burst number during the light phase with no change in light phase burst size. This suggests that reduced satiety signaling in the light phase underlies the change in water intake timing. Furthermore, the water intake that remained in the dark phase was associated with a reduction in burst size. This suggests that drinking in the absence of food has reduced hedonic value. When water intake was restricted to the light phase, burst number increased, again suggesting reduced satiety signaling under this manipulation. Burst size during WRL did not change in size and was similar to burst sizes taken during the BL light phase. This suggests that hedonic value does not contribute to the increased water intake in the light phase when access is restricted to this period, and in fact is reduced compared to dark phase intake under *ab libitum* conditions. This comparative reduction in burst size could be related to the differences in food intake during these times (more dark phase food intake in BL vs WRL). Together, both experimental manipulations suggest that altering water intake, either by shifting food intake or shifting water access, changes lick patterns via alterations in satiety signaling and that water intake in the absence of food, or with reduced food intake, is associated with a reduction in the hedonic value of the water in both males and females.

Despite food access having a large effect on water intake patterns, biological rhythms still play an important role in this behavior. As is the case with food intake [11], circadian patterns of water intake persist under constant dark or constant light conditions [10]. Light entrains most biological rhythms, and indeed blinding rats increase water and food intake in the light phase [1, 41]. Lesions to the pituitary gland, but not the inferior accessory optic tracts, primary optic tracts, or pineal gland, disrupt circadian patterns of water intake in rats, suggesting neuroendocrine involvement [41]. Furthermore, lesions to the suprachiasmatic nucleus, the central circadian clock, abolishes circadian water intake patterns, likely through direct projections to the hypothalamus [12, 13]. Given the circadian control of food intake, the strong association of water intake with food intake, and the inability to measure water intake for prolonged periods in the absence of food, disentangling the contributions of food and circadian rhythms on water intake patterns remains a challenge and requires additional research.

This series of experiments provides addition support for past studies demonstrating that water intake shifts to coincide with food access, even during prolonged periods of restricted food access [1, 3, 6, 16]. The results are the first to demonstrate that the shifted water intake is mediated by changes in satiety signaling, as increased burst number for water was associated with the change in water intake patterns [34]. These results are also the first to demonstrate that there are no sex differences in these intake patterns, beyond males consuming more food and water than females. Finally, while some food intake shifted into the light phase to coincide with water access, there was not a complete reversal of the circadian intake patterns, as observed with water intake when food was restricted to the light phase. Together this suggests that while both food and water influence intake of the other, eating has a greater effect on water intake patterns than drinking has on food intake patterns.

## Acknowledgements

We thank Kennedy Lamb, Grace Young, and Lasni Nishshanke for their technical assistance. This work was supported by National Science Foundation Awards 2019346 to JS and a Neuroscience Research Priority Areas (NRPA) Summer Undergraduate Research Award (SURA) to ELM.

## References

1. Zucker, I., Light-dark rhythms in rat eating and drinking behavior. Physiol Behav, 1971. 6(2): p. 115–26.

2. Siegel, P.S. and H.L. Stuckey, The diurnal course of water and food intake in the normal mature rat. J Comp Physiol Psychol, 1947. 40(5): p. 365–70.

3. Panksepp, J. and K. Krost, Modification of diurnal feeding patterns by palatability. Physiol Behav, 1975. 15(6): p. 673–7.

4. Greenwood, K., S. Armstrong, and G. Coleman, Persistence of rat nocturnal feeding and drinking during diurnal presentation of palatable diet. Physiol Behav, 1980. 24(6): p. 1119–23.

5. Spiteri, N.J., Circadian patterning of feeding, drinking and activity during diurnal food access in rats. Physiol Behav, 1982. 28(1): p. 139–47.

6. Fitzsimons, T.J. and J. Le Magnen, Eating as a regulatory control of drinking in the rat. J Comp Physiol Psychol, 1969. 67(3): p. 273–83.

7. Velasco Plaza, A., G.G. T, and M.T. Cachero, Circadian rhythms of food and water intake and urine excretion in diabetic rats. Physiol Behav, 1993. 54(4): p. 665–70.

8. Lensu, S., Tiitanen, P., Pohjanvirta, R., Circadian differences between two rat strains in their feeding and drinking micro-and macrostructures. Biological Rhythm Research, 2011. 42(5): p. 385–405.

9. Glendinning, J.I. and J.C. Smith, Consistency of meal patterns in laboratory rats. Physiol Behav, 1994. 56(1): p. 7–16.

10. Glantz, R.M., Circadian Rhythms in the Albino Rat: Effect of Illumination on Urine Excretion, Water Intake, and the Formation of the Anti-Diuretic Hormone. Physiol Behav, 1967. 2: p. 49–54.

11. Stoynev, A.G. and O.C. Ikonomov, Effect of constant light and darkness on the circadian rhythms in rats: I. Food and water intake, urine output and electrolyte excretion. Acta Physiol Pharmacol Bulg, 1983. 9(3): p. 58–64.

12. Stephan, F.K. and I. Zucker, Circadian rhythms in drinking behavior and locomotor activity of rats are eliminated by hypothalamic lesions. Proc Natl Acad Sci U S A, 1972. 69(6): p. 1583–6.

13. Van den Pol, A.N. and T. Powley, A fine-grained anatomical analysis of the role of the rat suprachiasmatic nucleus in circadian rhythms of feeding and drinking. Brain Res, 1979. 160(2): p. 307–26.

14. Kissileff, H.R., Food-associated drinking in the rat. J Comp Physiol Psychol, 1969. 67(3): p. 284–300.

15. Strominger, J.L., The relation between water intake and food intake in normal rats and in rats with hypothalamic hyperphasia. Yale J Biol Med, 1947. 19(3): p. 279–88.

16. Kutscher, C.L., Species differences in the interaction of feeding and drinking. Ann N Y Acad Sci, 1969. 157(2): p. 539–52.

17. Santollo, J. and D. Daniels, Fluid transitions. Neuropharmacology, 2024. 256: p. 110009.

18. Naneix, F., K.Z. Peters, and J.E. McCutcheon, Investigating the Effect of Physiological Need States on Palatability and Motivation Using Microstructural Analysis of Licking. Neuroscience, 2020. 447: p. 155–166.

19. Davis, J.D., The microstructure of ingestive behavior. Ann N Y Acad Sci, 1989. 575: p. 106–19; discussion 120-1.

20. Santollo, J., et al., Sex Differences in Salt Appetite: Perspectives from Animal Models and Human Studies. Nutrients, 2023. 15(1).

21. Santollo, J., Sex differences in AngII-Stimulated fluid intake. Exp Physiol, 2017.

22. Santollo, J. and D. Daniels, Multiple estrogen receptor subtypes influence ingestive behavior in female rodents. Physiol Behav, 2015. 152(Pt B): p. 431–7.

23. Asarian, L. and N. Geary, Sex differences in the physiology of eating. Am J Physiol Regul Integr Comp Physiol, 2013. 305(11): p. R1215–67.

24. Fukushima, A., et al., Sex differences in feeding behavior in rats: the relationship with neuronal activation in the hypothalamus. Front Neurosci, 2015. 9: p. 88.

25. Lok, R., J. Qian, and S.L. Chellappa, Sex differences in sleep, circadian rhythms, and metabolism: Implications for precision medicine. Sleep Med Rev, 2024. 75: p. 101926.

26. Williams, C.T., et al., Sex-Differences in Phenology: A Tinbergian Perspective. Integr Comp Biol, 2022. 62(4): p. 980–997.

27. Walton, J.C., J.R. Bumgarner, and R.J. Nelson, Sex Differences in Circadian Rhythms. Cold Spring Harb Perspect Biol, 2022. 14(7).

28. Santollo, J., A. Marshall, and D. Daniels, Activation of membrane-associated estrogen receptors decreases food and water intake in ovariectomized rats. Endocrinology, 2013. 154(1): p. 320–9.

29. Sortman, B.W., et al., Drinking microstructure analysis of direct controls of water intake in male and female rats. Physiol Behav, 2026. 310: p. 115314.

30. Demieri, E., et al., A microstructural analysis of sodium and water intake in male and female rats. Physiol Behav, 2026. 317: p. 115507.

31. Agostinelli, C.a.L., U., R package ‘circular’: Circular Statistics (version 0.5-2). 2025.

32. Gizowski, C., C. Zaelzer, and C.W. Bourque, Clock-driven vasopressin neurotransmission mediates anticipatory thirst prior to sleep. Nature, 2016. 537(7622): p. 685-8.

33. Johnson, R.F., et al., Investigations on the physiological controls of water and saline intake in C57BL/6 mice. Am J Physiol Regul Integr Comp Physiol, 2003. 285(2): p. R394–403.

34. Davis, J.D., G.P. Smith, and B. Singh, A microstructural analysis of the control of water and isotonic saline ingestion by postingestional stimulation. Physiol Behav, 1999. 66(3): p. 543–8.

35. Fitzsimons, J.T. and B.J. Simons, The effect on drinking in the rat of intravenous infusion of angiotensin, given alone or in combination with other stimuli of thirst. J Physiol, 1969. 203(1): p. 45–57.

36. Smith, G.P., John Davis and the meanings of licking. Appetite, 2001. 36(1): p. 84–92.

37. Davis, J.D. and G.P. Smith, Analysis of the microstructure of the rhythmic tongue movements of rats ingesting maltose and sucrose solutions. Behav Neurosci, 1992. 106(1): p. 217–28.

38. Spector, A.C., P.A. Klumpp, and J.M. Kaplan, Analytical issues in the evaluation of food deprivation and sucrose concentration effects on the microstructure of licking behavior in the rat. Behav Neurosci, 1998. 112(3): p. 678–94.

39. Davis, J.D. and M.C. Perez, Food deprivation-and palatability-induced microstructural changes in ingestive behavior. Am J Physiol, 1993. 264(1 Pt 2): p. R97-103.

40. Hsiao, S. and R.J. Fan, Additivity of taste-specific effects of sucrose and quinine: microstructural analysis of ingestive behavior in rats. Behav Neurosci, 1993. 107(2): p. 317–26.

41. Stephan, F.K. and I. Zucker, Rat drinking rhythms: central visual pathways and endocrine factors mediating responsiveness to environmental illumination. Physiol Behav, 1972. 8(2): p. 315–26.

